# Autotoxin-mediated voluntary triage in starved yeast community

**DOI:** 10.1101/2020.10.17.344093

**Authors:** Arisa H. Oda, Miki Tamura, Kunihiko Kaneko, Kunihiro Ohta, Tetsuhiro S. Hatakeyama

## Abstract

When organisms face crises, such as starvation, every individual should adapt to environmental changes (1, 2), or the community alters their behaviour (3–5). Because a stressful environment reduces the carrying capacity (6), the population size of unicellular organisms shrinks in such conditions (7, 8). However, the uniform stress response of the cell community may lead to overall extinction or severely damage their entire fitness. How microbial communities accommodate this dilemma remains poorly understood. Here, we demonstrate an elaborate strategy of the yeast community against glucose starvation, named the voluntary triage. During starvation, yeast cells release some autotoxins, such as leucic acid and L-2keto-3methylvalerate, which can even kill the cells producing them. Although it may look like mass suicide at first glance, cells use epigenetic “tags” to adapt to the autotoxin inheritably. If non-tagged latecomers, regardless of whether they are closely related, try to invade the habitat, autotoxins kill them and inhibit their growth, but the tagged cells can selectively survive. Phylogenetically distant fission and budding yeast (9) share this strategy using the same autotoxins, which implies that the universal system of voluntary triage may be relevant to the major evolutional transition from unicellular to multicellular organisms (10).

## Introduction

In a crisis, such as starvation, organisms should adapt at both individual (1, 2) and population levels (3–5). In unicellular organisms, the former has been intensively studied as an adaptation phenomenon (11–13), whereas the latter is poorly understood. Severe conditions decrease the carrying capacity (6), and unicellular organisms have to decrease the population size (7, 8). However, such an adjustment in cell number carries the risk of killing clonal cells. Thus, how cellular communities adapt to crisis without decreasing the fitness of clonal cells remains unknown.

Here, we report an elaborate survival strategy in crisis, named a voluntary triage. When the fission yeast, *Schizosaccharomyces pombe*, is cultured in glucose-limited conditions, where the carrying capacity is expected to decrease, cells release toxic molecules into the medium. Surprisingly, such a medium kills even the clonal cells of the toxin-producing cells when they are transferred from glucose-rich conditions. This may look like mass suicide at first glance. However, cells precultured in starved conditions continue to grow even in the conditioned medium, as they tag themselves to adapt to the toxins through starvation, and such adapted state is inheritable. In other words, cells autonomously differentiate into two types, adapted and non-adapted ones, and the cellular community selectively saves the former, just like triage in emergency medical care. Such voluntary triage works as a competitive strategy against both closely related and distant species. Starved yeast cells release toxins, which prevent an invasion of latecomers by killing them, as the Greek philosopher argued: the plank of Carneades (14). Surprisingly, the same strategy was seen in budding yeasts, which are phylogenetically distant relatives to the fission yeast (9). Indeed, we identified the same toxic molecules from both media conditioned by fission and budding yeast. Therefore, we hypothesised that the voluntary triage is evolutionarily conserved in fungal microbes.

## Results

To detect interactions in the population, we prepared conditioned media (CM) by culturing wild type (WT) fission yeast *S. pombe* for 30 h in the minimal media (MM) without glucose (0% MM) (see Fig. 1A for detailed procedure). We refer to this medium as the WT CM. When cells, which had been precultured in the MM with 3% glucose (3% MM), were cultured in the WT CM, they stopped growing for approximately 20 h and then resumed growing (see the red line in Fig. 1B). We termed this prolonged lag phase as the delay phase (see Supplementary Note 1 for measurement of the delay phase). If incubation time to prepare CM was longer than 15 h, such media also induced the delay phase, while shorter incubation time did not introduce such a phase (Fig. 1B). This indicated that in the early growth phase, cells released inhibitors for growth or depleted some of the nutrients required for such a phase.

**Fig. 1.**
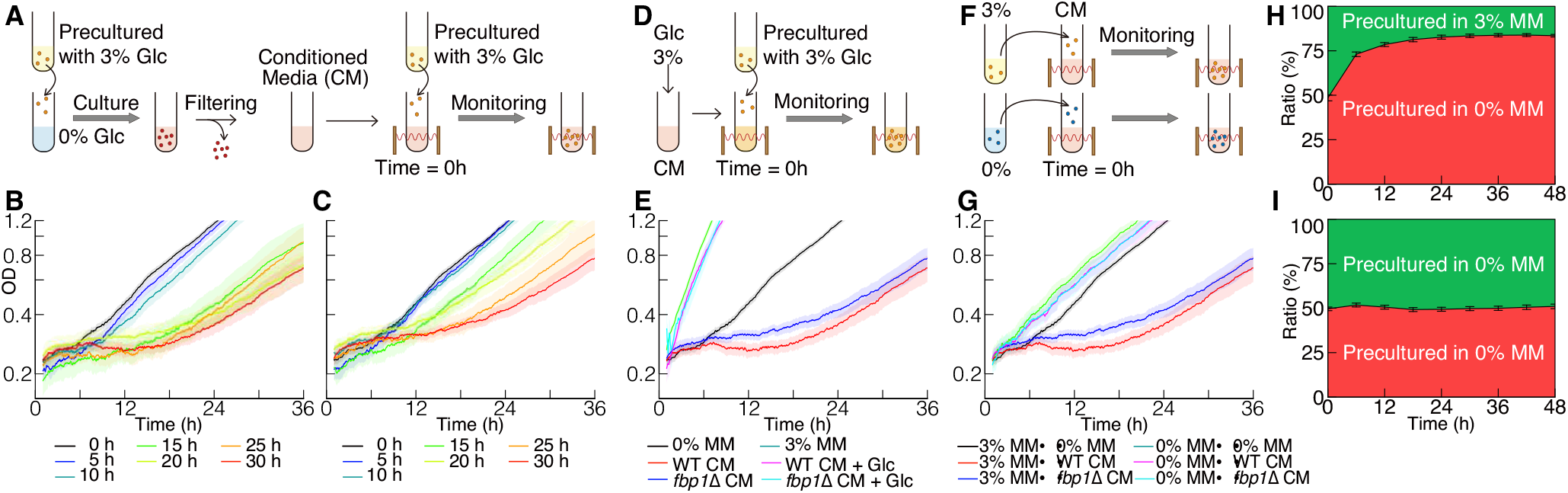
Conditioned media caused a delay phase during glucose starvation. A) Schematic illustration of the experimental procedure for (B) and (C). B and C) Growth curves of wild type (WT) cells in (B) WT- or (C) *fbp1*Δ cell-conditioned media (CM) without glucose. Different coloured lines indicate a moving average of optical density (OD) measured every minute in CM with different incubation times. Each line is an average of *n ≥* 10 samples, and the pale-coloured area indicates the standard error of the mean (SEM). D) Schematic illustration of the experimental procedure for (E). E) Growth curves of WT cells in CM with 3% glucose. Each line represents an average of *n ≥* 7 samples. F) Schematic illustration of the experimental procedure for (G). G) Growth curves of WT cells, precultured without glucose, in the CM. Each line represents an average of *n ≥* 7 samples. H and I) Competition assay in WT CM (H) between the cells precultured in 3 and 0% minimal media (MM) and (I) between the cells precultured in 0% MM. Green and red areas indicate the fraction of mNeonGreen- and mCherry-labelled cells, respectively, and overwriting outline characters indicate preculture conditions. Black vertical bars between two areas indicate SEM (number of each sample is 12).

To determine whether cells release inhibitors or deplete essential nutrients, we constructed a conditioned medium using a 1,6 bisphosphatase deletion mutant (*fbp1*Δ), which did not have a functional gluconeo-genetic pathway (15). Such a mutant strain could not grow without glucose (Fig. S1 and (16, 17)) and was expected not to consume the nutrients required for growth. The CM made using *fbp1*Δ cells (*fbp1*Δ CM) also caused the delay phase (Fig. 1C), as shown with the WT CM. This suggested that the delay phase resulted from the release of inhibitory molecules by cells rather than the depletion of nutrients.

In addition, when we administered a sufficient concentration of glucose to the CM (Fig. 1D) to recover the carrying capacity, cellular growth was not disrupted, and the delay phase was not observed, i.e., the growth curve of cells in such media was almost the same as that of those in MM with glucose (Fig. 1E). This indicated that inhibitory molecules in the CM worked only in the absence of glucose.

After the delay phase, the growth rate in the CM returned to almost the same level as in the MM. This suggested that the cells adapted to inhibitory molecules in the CM, and such an adapted state was expected to be inheritable. To verify the existence of the adapted state of cells, we precultured cells in the 0% MM and measured their growth in WT and *fbp1*Δ CM (Fig. 1F), and then no delay phase was observed (Fig. 1G). Furthermore, to verify whether adaptation to the inhibitory compound was due to genetic or epigenetic changes, we precultured cells that survived in the CM in unstarved condition, and again, cultured them in the CM; they showed a delay phase (Fig. S2). Moreover, we performed gDNA-seq of surviving cells and identified no unique SNPs and InDels, except for highly repetitive sequence loci, such as telomeres and centromeres (see Fig. S3 and Supplementary Note 2 for details). This indicated that the adapted cell was “tagged” epigenetically.

Plausible evolutionary significance of the release of inhibitory molecules and adapting to them is the inhibition of growth of different lineages of cells. When sugars around cells are depleted, they start to release inhibitory molecules while simultaneously adapting to such inhibitors. Then, the modified environment will inhibit the growth of latecomers, even if they are closely related. We performed a competition assay by artificially mimicking the above conditions; we simultaneously added cells that were precultured in glucose-rich and glucose-poor media into the CM with an equal amount in the beginning and observed their population dynamics (Figs. S4). Then, the fraction of adapted cells to unadapted cells continued to increase for 24 h and reached a steady-state (Figs. 1H and S5A), while the fifty-fifty ratio was maintained in the competition assay between adapted cells (Figs. 1I and S5B). In addition, the steady-state ratio of adapted and unadapted cells agreed with the ratio predicted from the growth curve observed in fresh and conditioned media in Fig. 1G (see also Supplementary Note 3). This implied that the combination of inhibitor release and adaptation caused population dynamics shown in Figs. 1H and I and selected the offspring of inhibitor-producing cells to survive.

The characteristics of the inhibitory molecules observed in the CM helped us isolate them. First, we identified chemical compounds in the freshly prepared MM as well as WT and *fbp1*Δ CM using capillary electrophoresis mass spectrometry (CE-MS). We identified 20 chemical compounds. From these candidates, we chose 12 chemicals that were included in both CM but not the fresh medium (see the yellow hatching region in the Venn diagram in Fig. 2A), because both CM initiated the delay phase. We further narrowed down 12 molecules by adding them to the MM according to the following criteria: 1) They did not change the growth rate significantly after the delay phase. 2) They had little effect on growth in the presence of glucose. 3) They caused a shorter delay phase in cells that had already adapted to starvation. Finally, we isolated two small molecules with similar structures: Leucic acid (HICA, Fig. 2B) and L-2Keto-3methylvalerate (2K3MVA, Fig. 2C). Note that some of the molecules in the candidate list were difficult to obtain commercially and could not be tested.

**Fig. 2.**
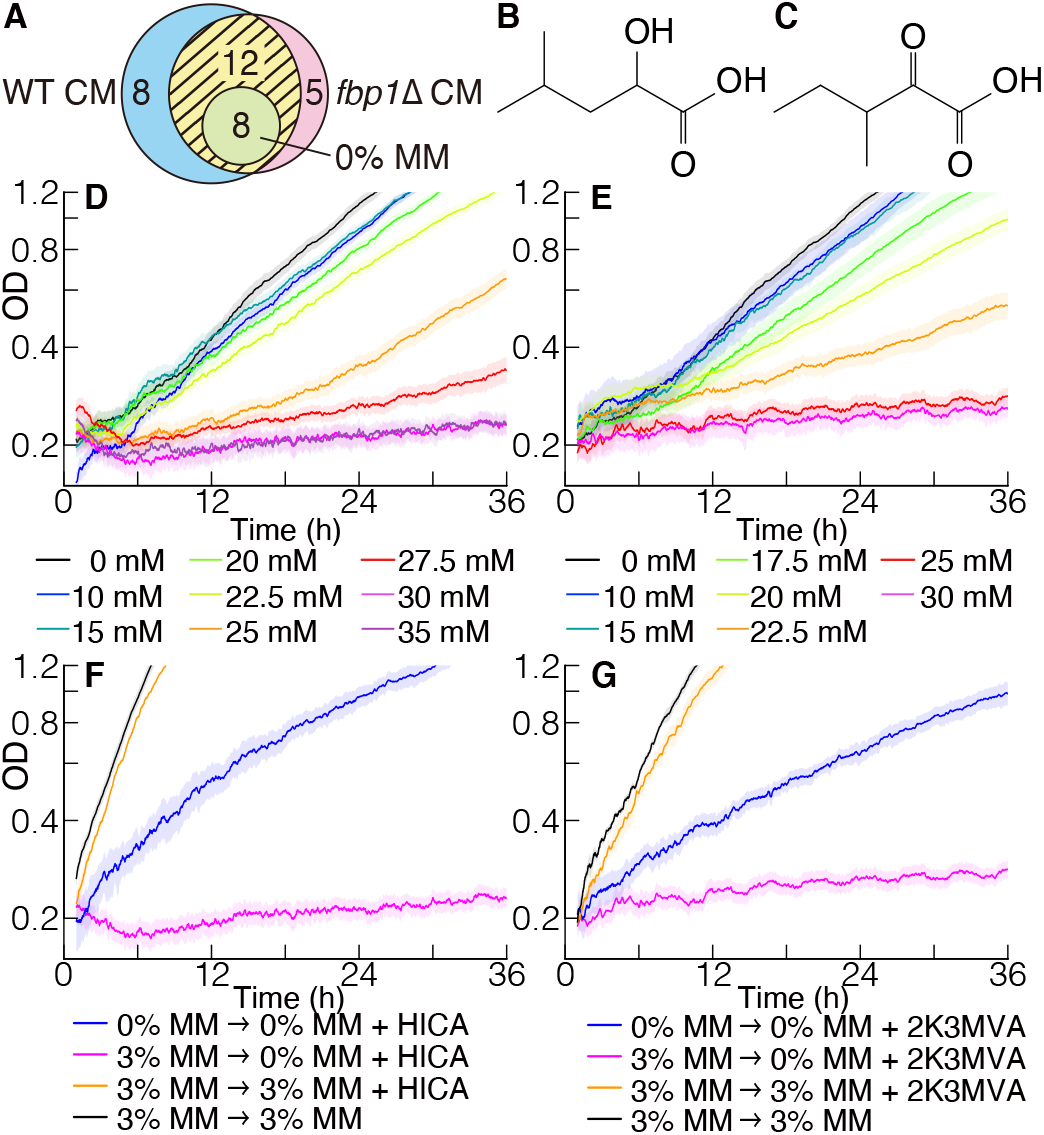
Identification of growth inhibitors. A) Venn diagram of compounds detected using capillary electrophoresis mass spectrometry (CE-MS). Compounds in 0% minimal media (MM), wild type (WT) conditioned media (CM), and *fbp1*Δ CM were analysed. Twenty compounds were detected in both WT and *fbp1*Δ CM, and eight of those were also detected in 0% MM. Thus, 12 compounds (yellow hatching area) were detected uniquely in both CM (see Supplementary Table 1 for details of the detected molecules). B and C) The structure of (B) HICA and (C) 2K3MVA. D and E) Growth curves in 0% MM in the presence of (D) HICA and (E) 2K3MVA. WT cells precultured in 3% MM were transferred to 0% MM with various concentrations of the inhibitory compound at 0 h. Each line represents the average of *n ≥* 6 samples. F and G) Effects of adaptation and glucose administration on growth curves in the presence of (F) 30 mM HICA or (G) 25 mM 2K3MVA. The blue line indicates the growth curve of WT cells precultured in 0% MM in 0% MM with the inhibitory compound. The orange line is a growth curve of WT cells in MM with the inhibitory compound and 3% glucose. The pink and black lines indicate growth curves in 0% MM with the inhibitory compound and 3% MM as controls, respectively. Each line represents an average of *n ≥* 4 samples.

The two inhibitory molecules had similar characteristics. When the concentration of these molecules was not sufficient, we never observed the delay phase (Figs. 2D and E). Then, the more we administrated the inhibitors, the longer the delay phase was. Finally, if the concentration was higher than the critical concentrations (30 mM for HICA and 25 mM for 2K3MVA), cell growth was thoroughly repressed. Notably, there are two optical isomers of HICA, both of which caused a growth delay at the same concentration (Fig. S6). When glucose was added to the MM simultaneously with inhibitory molecules, cell growth was not disrupted (see the orange line in Figs. 2F and G). Moreover, even under the administration of such high concentration where cells stopped growing, cells that had been adapted to glucose starvation grew (see blue line in Figs. 2F and G). These correspondences of inhibitory molecules with the CM implied that the release of HICA and 2K3MVA was one of the causes of voluntary triage during glucose starvation.

How do the inhibitory molecules cause the delay phase? There are two possible mechanisms: Delay of initiation of growth in each cell or death of the majority of cells. In the latter, the concentration of living cells is masked by that of dead cells in the OD measurement, and an apparent delay phase will be observed until the concentration of living cells exceeds that of dead cells red(see Supplementary Note 4 and Fig. S7). To verify which hypothesis is correct, we counted the number of dead cells by staining them with phloxine B, a red dye, often used to check yeast viability (18) (Figs. 3A–H). Then, over 80% of cells, which were cultured in the presence of a higher concentration of inhibitory molecules, were dyed in red (Figs. 3G and H). Indeed, the majority of cells during the delay phase were stained red and showed a typical rod shape, as observed under a microscope, which are characteristics of dead cells (19, 20) (Figs. 3E and F). In contrast, only a small number of cells showed a spherical shape similar to cells cultured in MM without glucose. This indicated that only a small portion of the cells survived and continued to divide in the presence of inhibitory molecules. Similarly, the death rate in WT and *fbp1*Δ CM increased (Figs. 3B, C, and G). Furthermore, cells precultured in the starved condition, which did not show a delay phase in the presence of the inhibitory molecules, were mostly alive (Fig. 3H). This suggested that HICA and 2K3MVA induced cell death, which was the primary cause of the delay phase (see also Supplementary Note 4, Figs. S8 and S9).

**Fig. 3.**
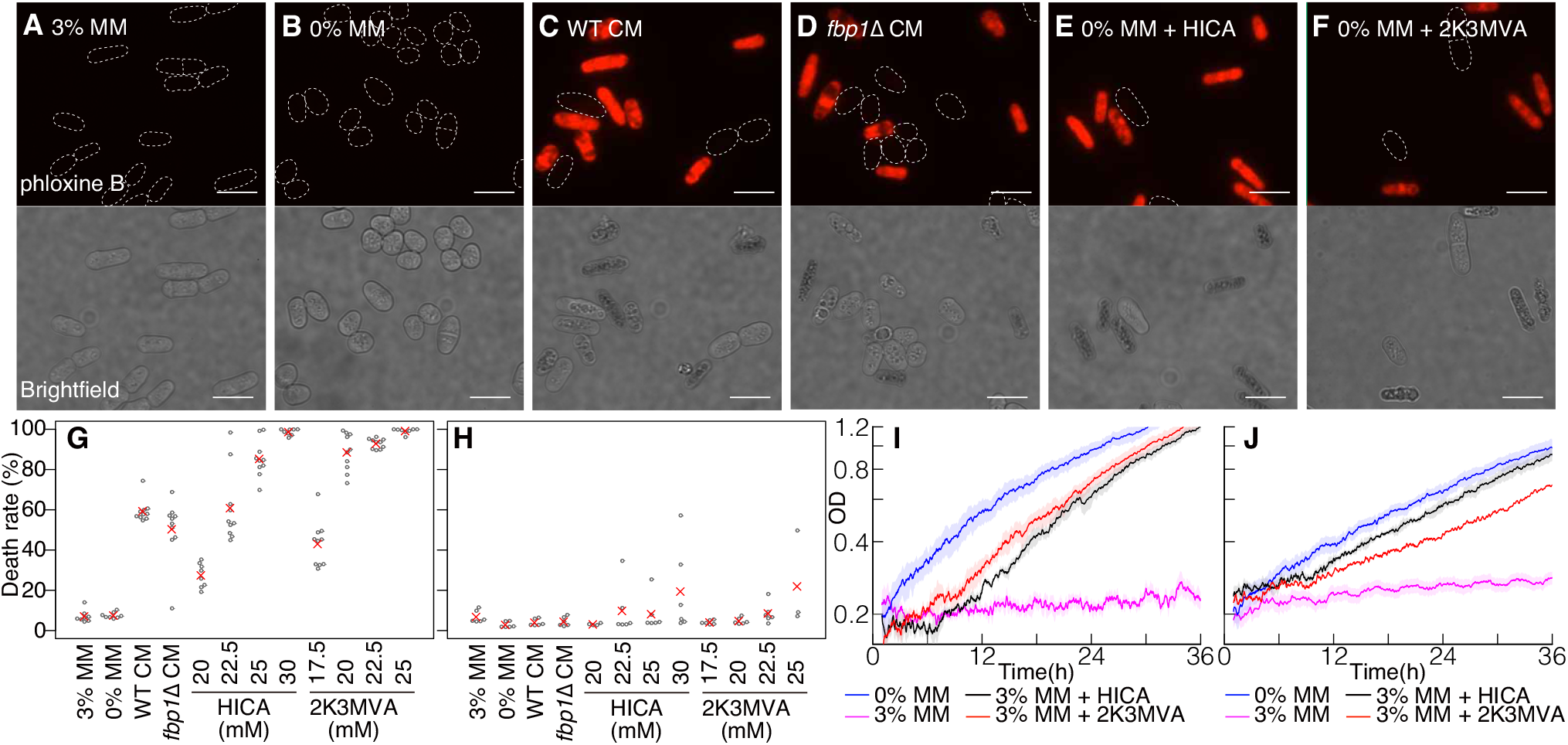
Identified molecules kill cells and also facilitate cell adaptation to the molecule. A–F) Fluorescent (upper) and brightfield (bottom) microscopic images of wild type (WT) cells in various media. Cells precultured in 3% minimal media (MM) were transferred to (A) 3% MM, (B) 0% MM, (C) WT conditioned media (CM), (D) *fbp1*Δ CM, (E) 0% MM with 25 mM HICA, and (F) 22.5 mM 2K3MVA. In fluorescent microscopic images, dead cells were stained with Phloxine B. Scale bar indicates 10 *μ*m. G and H) The dyed cell ratio after 8 h of incubation. Cells were precultured in (G) 3% MM (*n* = 8-10) or (H) 0% MM (*n* = 3-6). Grey dots represent the dyed cell ratio in each sample, and red crosses represent the mean value. I and J) Growth curves of cells precultured in the presence of one of the inhibitory molecules along with 3% glucose, in 0% MM with (I) 30 mM HICA or (J) 25 mM 2K3MVA. Cells were precultured in 0% MM (blue line), 3% MM (pink line), 3% MM with 30 mM HICA (black line), and 3% MM with 25 mM 2K3MVA (red line).

Note that the identified toxins also facilitate cell adaptation in a condition-dependent manner. When we precultured cells in 3% MM with an inhibitory molecule, HICA or 2K3MVA, they grew in 0% MM with a sufficiently high concentration of inhibitory molecules that stopped the growth of non-adapted cells (Figs. 3I and J). This indicated that the molecules we identified had two distinguishable effects: To kill starving cells and help non-staving cells to adapt. We also characterized the gene expression pattern in inhibitory molecule-adapted cells, and confirmed that the metabolic state of cells grown in the presence of both glucose and inhibitory molecules was different from that of those grown in glucose starvation and as close to that of those grown in 3% MM (see Supplementary Note 5 and Fig. S10).

Voluntary triage is not a unique characteristic of *S. pombe* but is widely observed in unicellular fungi. We cultured two strains of budding yeast *Saccharomyces cerevisiae*, which are phylogenetically distant from *S. pombe*, and prepared CM using them. Then, we found that such CM also initiated the delay phase in the growth of media producers (Figs. 4A and S11A). Moreover, we detected the same toxic molecules, HICA and 2K3MVA, in the media conditioned with *S. cerevisiae*. In addition, the administration of such toxins to a MM without glucose initiated the delay phase in a concentration-dependent manner (Figs. 4B, C, and S11B and C). Cells precultured in the starved condition did not show delay phase in their CM or 0% MM with an inhibitory molecule (Figs.4D, E, S11D, and E). This suggested that the same strategy with the same molecules, as observed in *S. pombe*, was evolutionarily conserved among distant species. In addition, media conditioned with distant species also initiated the delay phase (Figs. 4F, G, and H), i.e., media conditioned with *S. pombe* inhibited the growth of *S. cerevisiae* and vice versa. Therefore, such a strategy was universally effective from closer to distant species.

**Fig. 4.**
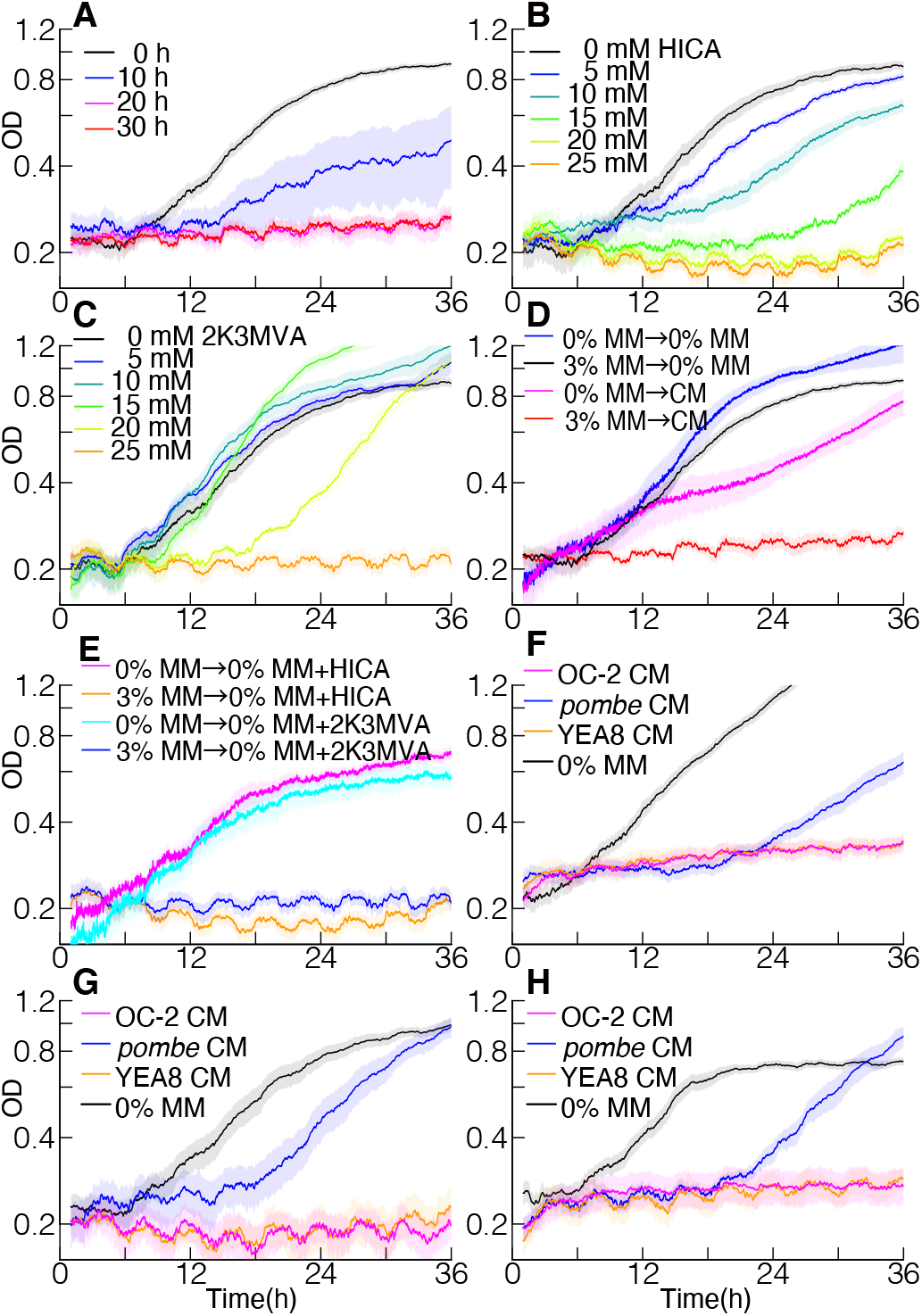
Media conditioned with various strains of yeasts initiated the delay phase. A) Growth curves of *S. cerevisiae* (OC-2) cells in media conditioned with themselves. Different coloured Lines indicate growth curves of conditioned media (CM) at different incubation times. Each line represents an average of *n ≥* 6 samples. B) Growth curves of OC-2 in 0% minimal media (MM) with various concentrations of HICA. Each line represents an average of *n ≥* 5 samples. C) Growth curves of OC-2 in 0% MM with various concentrations of 2K3MVA. Each line represents an average of *n ≥* 5 samples. D) Growth curves of OC-2 precultured in 0 or 3% MM in OC-2 CM. Each line represents an average of *n ≥* 6 samples. E) Growth curves of OC-2 precultured in 0 or 3% MM in 0% MM with 25 mM HICA or 25 mM 2K3MVA. Each line represents an average of *n ≥* 6 samples. F, G, and H) Growth curves of (F) *S. pombe*, (G) OC-2, and (H) YEA8 in media conditioned with *S. pombe*, OC-2, or YEA8 for 30 h. Each line represents an average of *n ≥* 6 samples.

## Discussion

In this paper, we reported a new ecological strategy for microbes; the voluntary triage. It seems similar to the toxin-antitoxin system, such as bacteriocins in bacteria (21, 22) and killer factors in yeast (23, 24) and paramoecium (25) but it is fundamentally different. Although the toxin-antitoxin system does not kill the clonal cells (26), the voluntary triage kills even the clonal cells if they have not been adapted to toxins. Such a strategy, appearing as a suicide at first glance, helps the yeast to select an appropriate offspring that produces toxins and selfishly purify their genome from closely related species. Moreover, voluntary triage overcomes the problems of the toxin-antitoxin system. In such a system, toxin producers should continuously produce antitoxins to protect themselves, and the maintenance of this state is a heavy burden for them (27). Thus, the toxin producer is lost to a cheater, which only has the antitoxin system, whereas the cheater loses to cells having neither any toxin nor immunity (28). In contrast, the voluntary triage does not cost much because the adaptation mechanism is usually offed without the toxin. This suggests that voluntary triage is resistant to cheaters.

We found that distant yeast species universally conserved voluntary triage, even at the molecular level. This might be because the toxins in the reported system are simple molecules, while toxins in the toxin-immunity systems in bacteria and yeast are highly evolved proteins. In the bacteriocin system of *Escherichia coli*, toxins and receptors set off an arms race between the diversification of toxins and enhancement of their recognition by modifying the structure of proteins (29). In contrast, in the voluntary triage, the targets of toxins are insensitive to the detailed structure of molecules and are not specific receptors. Indeed, the enantiomers of HICA have the same activity, where both D- and L-forms of HICA cause the delay phase at the same concentration (Fig. S6). Moreover, HICA was first identified from fermented products of a bacterium, *Lactobacillus plantarum* (30), and is toxic to various bacterial species and *Candida* and *Aspergillus* species (31). This suggests that HICA and 2K3MVA targets are universally conserved pathways. Therefore, the toxins we found were effective against a range of cells; from clones to distant species.

The voluntary triage we reported might play an important role in understanding the origin of multicellularity. Multicellular fungi fall into clades 8–11, while there are only four clades apart from fungi (32, 33), indicating that transitions from unicellular to multicellular and multicellular to unicellular organisms occur easily in fungi. To form a complex multicellular body, mutual activation of growth, as well as growth inhibition and programmed cell death pathway, are essential (34). Thus, the evolutionary origin or vestige of both mutual activation and inhibition of growth should be observed even in unicellular fungi. Indeed, multiple species of unicellular fungi have the former as quorum sensing (35). However, the latter has not been reported. The mechanism we found here meets the criteria required for growth inhibition for multicellularity (36, 37), that is, the toxins cause cell death depending on the cell state and smoothly diffuse from cell to cell. A recent artificial evolution experiment demonstrated that multicellular “snowflake” yeast repeatedly evolved 15 times from unicellular *S. cerevisiae* and showed apoptosis to keep the original size constant (38). This suggests that the origin or vestige of multicellularity is embedded in the unicellular yeast. The relationship between intercellular communication in unicellular cells and multicellularity is key to solving the enigma of major transitions in evolution (10), and our study provides a significant milestone.

## ACKNOWLEDGEMENTS

This work was partially supported by the Basic Science Research Projects of Sumitomo Foundation, the Ohsumi Frontier Science Foundation, and Japan Society for the Promotion of Science (JSPS) KAKENHI (19K16070) to A.O. The authors would like to thank H. Nakaoka for his help in preparing *S. pombe* strains.

