## Supplementary Materials for "Autotoxin-mediated voluntary triage in starved yeast community"

#### Materials and Methods

**Continuous measurement of growth.** Continuous measurement of the optical density (OD) of yeast cultures was performed using ODBox-C/ODMonitor systems (Taitec Corp., Saitama, Japan). Such an apparatus detects the rate of transmitted light (950 nm) of shaken liquids in test tubes every minute. The raw data of transmitted light intensity was linearly converted into OD at 600 nm ($OD_{600}$) using the measurement data via iMark Microplate Absorbance Reader (Bio-Rad Laboratories inc., Hercules, CA, USA). Detailed delay phase and death rate analyses are described in Supplementary Note 1 and 4.

**Yeast strains and culture conditions.** Haploid strains, L972 ($h^-$ 972) and AR110 ($h^-$ *fbp1*Δ) were used for the *S. pombe* experiments. Wine yeast OC-2 (IAM4274) and baker's yeast YEA8 were used for the *S. cerevisiae* experiments. Strain information is also listed in Supplementary Table 3. Initially, cells were precultured at 30°C in yeast extract with supplements (YES; 3% Glucose, 0.5% yeast extract, 200 mg/L Adenine, 100 mg/L Uracil, 200 mg/L Histidine, 200 mg/L Leucine) liquid media. Then, cells were transferred to liquid minimal mediua (MM) with 3% glycerol and 3% glucose and precultured at 30°C for about 24 h until $OD_{600}$ reached to 0.15-0.2 unless otherwise noted. The final concentration of MM was as follows; 14.7 mM potassium hydrogen phthalate, 15.5 mM $Na_2HPO_4$, 93.5 mM $NH_4Cl$, 0.05% EtOH, 5.2 mM $MgCl_2$, 0.1 mM $CaCl_2 \cdot H_2O$, 13.4 mM KCl, 0.28 mM $Na_2SO_4$, 4.2 $\mu$M pantothenic acid, 81.2 $\mu$M nicotinic acid, 55.5 $\mu$M inositol, 40.8 nM D-Biotin, 4.76 $\mu$M citric acid, 8.09 $\mu$M boric acid, 2.37 $\mu$M $MnSO_4$, 1.39 $\mu$M $ZnSO_4{\cdot}7H_2O$, 0.74 $\mu$M $FeCl_3{\cdot}6H_2O$, 0.81 $\mu$M $(NH_4)_6Mo_7O_{24} \cdot 4H_2O$, 0.60 $\mu$M potassium iodide, 0.16 $\mu$M $CuSO_4 \cdot 5H_2O$ 200 mg/L adenine, 100 mg/L uracil, 200 mg/L histidine, 200 mg/L leucine. The culture was then inoculated into various media including MM with 3% glycerol without glucose (0% MM) and conditioned media. Before cells were transferred, they were washed twice with the new media.

**Preparation of conditioned media.** Cells were precultured in 3% MM and transferred to 0% MM and cultured. The supernatants were then filtered using a 0.22 $\mu$m PVDF membrane (Millex Sterile Filter Unit, Merck Millipore, Burlington, MA, USA).

DRAFT

**Flow cytometry analysis for competition assays.** Cells were genetically tagged with mNeonGreen (HN98) or mCherry (HN101). Strain information is also listed in Supplementary Table 3. They were precultured in 3% or 0% MM and transferred to 0% MM and WT CM.

mCherry and mNeonGreen fluorescence were detected at 561 nm and 488 nm and collected with 615/20 and 530/30 emission filters of NovoCyte flow cytometer (ACEA Biosciences, Inc., San Diego, CA, USA), respectively. mNeonGreen-tagged strain was constructed from pFA6a-mNeonGreen-HIS3MX6, which was a gift from Wei-Lih Lee (Addgene plasmid #129100 ; http://n2t.net/addgene:129100 ; RRID:Addgene_129100) The gating strategy for competition assay is shown in Supplementary Fig. 4.

**Capillary electrophoresis-time-of-flight mass spectrometry.** Media composition was measured using an Agilent Capillary Electrophoresis System, and data were processed by Human Metabolome Technologies Inc. (Turuoka, Japan) in anion and cation analysis modes according to a previously published method (1). Peaks with signal/noise ratios of more than 3-fold were extracted using MasterHands software ver2.17.1.11 (Keio University, Turuoka, Japan), and HMT's metabolite libraries were mined, including more than 900 metabolites to search for metabolites in the media.

**Search for growth inhibitors.** Capillary electrophoresis-time-of-flight mass spectrometry (CE-MS) was performed in two biological replicates for 0% MM, and *fbp1*Δ CM, and in three biological replicates for WT CM. We extracted compounds found in both samples (compounds found in only one sample are described as N.R. in Supplementary Table 1), and 12 candidates of growth inhibitors were identified following the criteria described in the main text. Every detected compound was added to freshly prepared 0% and 3% MM at a final concentration of 10–40 mM. Then, the growth rate of cells in such media was measured using ODBox-C/ODMonitor systems.

**RNA extraction and gene expression analysis using reverse transcriptase-polymerase chain reaction.** For reverse transcriptase-polymerase chain reaction (RT-PCR) analysis, three biological replicates were measured with two technical replicates for each sample. 15–20 mL cultures 24 h after media change were prepared as described above. Cells were pelleted down and frozen in liquid nitrogen. Frozen cell pellets were resuspended in 250 $\mu$L beads buffer (75 mM $NH_4OAc$, 10 mM EDTA, pH 8.0) at 65 °C with 200 $\mu$L acid-washed glass beads (Sigma), 25 $\mu$L of 10% SDS and 300 $\mu$L of acid-phenol:chloroform (pH4.5, Thermo Fisher Scientific Inc.). The samples were vortexed three times 1 min each at 1 min intervals and incubated at 65 °C, followed by a 10 min incubation at 65 °C, 1 min vortexing, and 15 min centrifugation at room temperature (16,000 × g). Then, the upper aqueous phase was transferred to a fresh tube with 200 $\mu$L of beads buffer and 400 $\mu$L of phenol/chloroform/isoamyl alcohol (25:24:1, Sigma). Tubes were vortexed briefly and centrifuged at 4°C, 16,000 × g for 15 min. The upper aqueous phase was transferred to a fresh tube with 600 $\mu$L ice-cold isopropanol and 21 $\mu$L 7.5 M $NH_4OAc$. Tubes were vortexed briefly and centrifuged at 4°C, 16,000 × g for 30 min. After removing the supernatant, the pellet was washed with 70% ethanol, air-dried, and resuspended in nuclease-free water.

For reverse transcription, PrimeScriptRT Reagent Kit with gDNA eraser (Takara Bio Inc., Japan) was used following the manufacturer's instructions. The cDNA libraries were quantified using StepOne Real-Time PCR system (Thermo Fisher Scientific Inc.) with the KAPA SYBR FAST qPCR Master Mix (2X) Kit (Sigma-Aldrich, St. Louis, MO, USA) following the manufacturer's instructions. PCR primers used for qPCR are listed in Supplemental Table 4.

**Flow cytometry analysis to measure the dead cell ratio.** Phloxine B (final concentration of 10 $\mu$g/ml, Sigma Aldrich) was used to dye dead cells. The fluorescent signal from phloxine B was detected at a wavelength of 488 nm and collected using 695/40 and 586/20 emission filters on a NovoCyte flow cytometer (ACEA Biosciences, Inc., San Diego, CA, USA). The gating strategy for the identification of dead cell population is shown in Supplementary Fig. 9.

**Microscopy and imaging analysis.** All images were captured using a microscope equipped with UPlanSApo 100x/1.40 Oil Objective (Olympus, Tokyo, Japan) on an EVOS FL Imaging System (Thermo Fisher Scientific Inc., Waltham, MA, USA). Images were acquired using a SonyICX285AL monochrome CCD camera controlled with built-in software for image acquisition. Fluorescence signal from phloxine B was detected at a wavelength of 530 nm and collected by EVOS Light Cube (Texas Red), which contains a 628/32 emission filter (Thermo Fisher Scientific Inc.).

**Measurement of surviving cell growth in conditioned media.** WT cells were cultured in WT CM for 24 h, and the culture was spread on YES agar plates to isolate single colonies. After incubating at 30 °C for three days, 24 colonies were picked and cultured in YES liquid medium for 24 h and then in 3% MM for 24 h. Then, the cells were cultured in WT CM again, and their growth curves were constructed using ODBox-C/ODMonitor systems.

**Genome sequencing of surviving cells in conditioned media.** WT cells were cultured in WT CM with phloxine B (final concentration of 10 $\mu$g/ml) for 24 h. Unstained living cells were selected using a cell sorter SH-800 (SONY) using a 488 nm excitation wavelength laser with a 525/50 emission filter and 561 nm excitation wavelength laser with a 617/30 emission filter. Then, they were cultured in YES liquid medium at 30°C for three days and pelleted down to extract genomic DNA. The cell pellet was resuspended in 150 $\mu$l of STES Buffer (0.5 M NaCl, 0.01 M EDTA, 1% SDS) with 150 $\mu$l of glass beads and 150 $\mu$l of phenol-chloroform. The samples were vortexed for 10 min, followed by a 10-min centrifugation at room temperature, 16,000 × g. The upper aqueous phase was transferred to a fresh tube with 150 $\mu$l of phenol-chloroform. The tubes were vortexed, followed by a 10-min centrifugation at 16,000 × g at room temperature. The upper aqueous phase was transferred to a fresh tube and mixed with 15 $\mu$l of 3 M NaOAc and 375 $\mu$l of ethanol. The samples were mixed briefly and centrifuged for 10 min at room temperature (16,000 × g). After the supernatant was removed, the pellet was rinsed with 70% ethanol, air-dried, and resuspended in nuclease-free water.

cDNA libraries were prepared using the NEBNext Ultra II DNA Library Prep Kit for Illumina (New England Biolabs, Inc. Ipswich, MA, USA), following the manufacturer's instructions. cDNA libraries were sequenced using the Illumina paired-end technology on MiSeq with MiSeq Reagent Kit v3 (600-cycle; Illumina, San Diego, CA, USA). Detailed information is provided in Supplementary Note 2. Sequences were deposited in DDBJ with DRA BioProject Acession Number: PRJDB10422.

DRAFT

#### Supplementary Note 1: Definition of the length of delay phase and growth rate

In general, when we transfer cells from a glucose-rich medium to a glucose-poor medium, cells show a lag phase where they switch intracellular activities to adapt to the new environment before starting growth. If we transferred cells to conditioned media and media with inhibitory molecules, a delay phase was observed in addition to the lag phase. We defined the length of lag and delay phases as $\tau$. We assumed that the lag and delay phases were linearly separable, and thus a difference between $\tau$ and $\tau_0$, which was $\tau$ for 0% MM, would give the length of the delay phase. Here, $\tau$ was measured as the time when the OD reached $2a_0$, where $a_0$ is the OD at the initial state. It was measured as $a_0$ as the average OD from 1–2 h because OD fluctuates in time, especially during the 1st hour, owing to the apparatus.

In addition, we defined the steady-state growth rate of cells after the lag and delay phases as $r$ (see also Supplementary Fig. 7 for the definition of $\tau$, $a_0$, and $r$). $r$ was measured within different ranges in different media due to variations in the delay phase length; 20–25 h for 0% MM, 25–35 h for WT CM, from 35–50 h for *fbp1*Δ CM, 15–30 h for 20 mM HIAC, 22.5 mM HICA, and 17.5 mM 2K3MVA, 30–45 h for 25 mM HICA, 25–40 h for 20 mM 2K3MVA, and 30–60 h for 22.5 mM 2K3MVA.

See Supplementary Table 2 for the measured $\tau$, $r$, and $a_0$.

#### Supplementary Note 2: Identification of mutated sites in surviving cells in conditioned media

We obtained $1.2 \times 10^6$ read pairs or more for each sample. After the removal of low-quality reads (phred quality $\geq$ Q15 and $>$ 10 base limit) using fastp (an ultra-fast all-in-one FASTQ pre-processor; version 0.20.0, (2)), we aligned the Illmina short reads to the reference genome of *Schizosaccharomyces pombe* (version 2018.09.04 (3)) on the PomBase database ((4)) using BWA (version 0.7.7, (5)). The average coverage depth for the reference chromosomes was 90× or more for each sample. We analysed mutations in each aligned dataset using the Genome Analysis Tool Kit-HaplotypeCaller software (GATK; version 4.1.4.1, (6, 7)) with the default parameters with an option of 'ploidy=1'. Then, we refined the raw variants identified by HaplotypeCaller using Variant filtration, with parameters 'QD < 2.0 or FS > 60.0, MQ < 40.0, or MQRankSum < -12.5 or ReadPosRankSum < -8.0 or SOR > 4.0', as recommended. We identified 271 and 268 SNPs and InDels, respectively, in two replicates of WT samples. Also 285 and 292 SNPs and InDels, respectively in the surviving samples. We searched for "common" and "unique" mutations in each sample using vcftools (version 0.1.13, (8)).

We classified each mutation into three groups: 1) Mutations that are common in all four samples; 2) mutations detected in both original WT and survivor samples but not in all samples; 3) mutations detected uniquely in a particular sample. We plotted the quality scores of mutations in each group as histograms in Supplementary Fig.3.

Quality scores of mutations commonly found in all four samples (yellow bars in Supplementary Fig. 3) were relatively high, and such mutations in the group 1 were expected to be the original SNPs and InDels in WT and be not the result of culturing in WT CM.

Mutations in group 2 tended to show lower quality scores than those in group 1 (see red bars in Supplementary Fig. 3). Mutations that were detected in only one of the survivor samples or both WT replicates, and one of the two survivor samples might have resulted from culturing in WT CM. Therefore, these 35 mutations were checked, and mutations were false positives.

Mutations in group 3 were unique mutations found in one sample (see blue bars for WT and cyan bars for surviving cells in Supplementary Fig. 3). To check whether these mutations arose during the culture in WT CM or not, we analysed all 84 unique mutations in the surviving samples. These unique mutations were located at repetitive sequences around telomeres and centromeres, where false positives are often found (9). In addition, the quality scores for these mutations were the lowest among the three groups; therefore, they were false positives.

DRAFT

#### Supplementary Note 3: Estimation of the steady-state ratio of cells in the competition assay

In the competition assay, adapted cells started to grow earlier than unadapted cells because they showed neither the lag nor the delay phase. Later, unadapted cells adapted to toxins, and their growth rate was the same as adapted cells. Hence, the ratio of adapted to unadapted cells reached a constant value at the steady-state. We calculated the steady-state ratio of cells as follows: The adapted cell growth rate was $r$ during $\tau$ hours, while the concentration of unadapted cells doubled after $\tau$ hours. Hence, if we assumed that the ratio of cells reached the steady state immediately after $\tau$ hours, the steady-state fraction of the adapted cells would be given as

$$\frac{\exp(r\tau)}{\exp(r\tau)+2} = \frac{\exp(0.056 \times 32.77)}{\exp(0.056 \times 32.77)+2} = 0.758. \quad \textbf{(1)}$$

This estimated value agreed well with the measured values obtained in the competition assay (see Fig. 2 in the main text).

#### Supplementary Note 4: Estimation of the death rate from growth rate and the length of delay phase

We assumed that cells were divided into two types: dead cells, which did not grow, and living cells, which showed exponential growth. The summation of both cells was observed as OD (see Supplementary Fig. 7). Then, the growth curve was drawn as follows:

$$\mathrm{OD}(t) = da_0 + (1-d)a_0 \exp(rt), \quad \textbf{(2)}$$

where $d$ is the death rate at time $= 0$. Thus, $d$ is calculated from $r$ and $\tau$ as follows:

$$d = \frac{2-\exp(r\tau)}{1-\exp(r\tau)}, \quad \textbf{(3)}$$

We estimated the death rate for each sample. Estimated death rates were well correlated with flow cytometry results (see Supplementary Fig. 8).
In 0% MM and CM, the estimated values were slightly higher than the measured values. In Eq. 2, we ignored the lag phase of living cells, which might cause an overestimation of the death rate. Indeed, the difference between the estimated and measured death rates in CM as almost the same as that in 0% MM.
In addition, at a high concentration of 2K3MVA, death rates were underestimated. In Eq. 2, we assumed that the growth rate was independent of time. If the growth rate decreased with time; that is, the concentration of living cells was given as a convex function, the death rate would be underestimated. In media with a high concentration of 2K3MVA, $r$ was much lower than that in other media (see Supplementary Table 2). This suggested that administration of 2K3MVA at a high concentration decreased the growth rate over time. Then, the death rate was underestimated. In any case, the delay phase was mainly caused by cell death.

#### Supplementary Note 5: Metabolic states of cells adapted to inhibitory molecules.

We measured gene expression in cells grown in the presence of both glucose and inhibitory molecules.
During growth without glucose, cells adapted to both glucose starvation and inhibitory molecules. These two adapted states seemed to be inseparable at first glance: Cells autonomously adapted to the inhibitory molecules when they adapted to glucose starvation, and vice versa. It is impossible to verify whether the two adaptations were inseparable or not via experiments using the CM, because two types of stresses, glucose starvation and inhibitory molecules, always coexist and are inseparable. Thus, we separated these two stresses with the aid of isolated molecules.
By comparing the gene expression of the genes involved in the metabolism of cells grown under glucose starvation, or with inhibitory molecules but not under glucose starvation, we found that the latter was distinct from the former and similar to that in 3% MM (Fig. S10). In 0% MM, cells took up glycerol and produced glucose via gluconeogenesis. Thus, the expression of genes involved in glycerol transportation (*gld1*, *dak1/2*, *fbp1*) were upregulated. With glucose and inhibitory molecules, however, cells did not express these genes, while with 2K3MVA, glycerol transporter and *dak1/2* expressions were upregulated. Further, although the expression of genes involved in the glycolysis pathway, from *fba1* to *pyk1*, were downregulated during glucose starvation, they were maintained in cells grown in the presence of glucose and inhibitory molecules. This suggested that adaptation to inhibitory molecules did not require changes in glycolysis and gluconeogenesis pathways and was different from adaptation to glucose starvation.

DRAFT

### Supplementary Figures

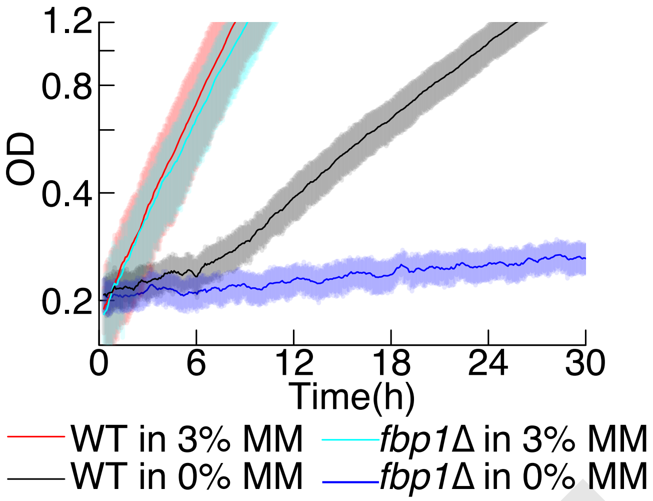


**Supplementary Fig. 1. Growth curve of *fbp1*Δ without glucose.** *fbp1*Δ and WT cells were precultured in 3% MM and then transferred to 0% or 3% MM. Growth curves of *fbp1*Δ cells in 0% and 3% are shown in blue and light blue, respectively, and those of WT cells are shown in black and red, respectively. Each line represents an average of 3-7 samples.

DRAFT

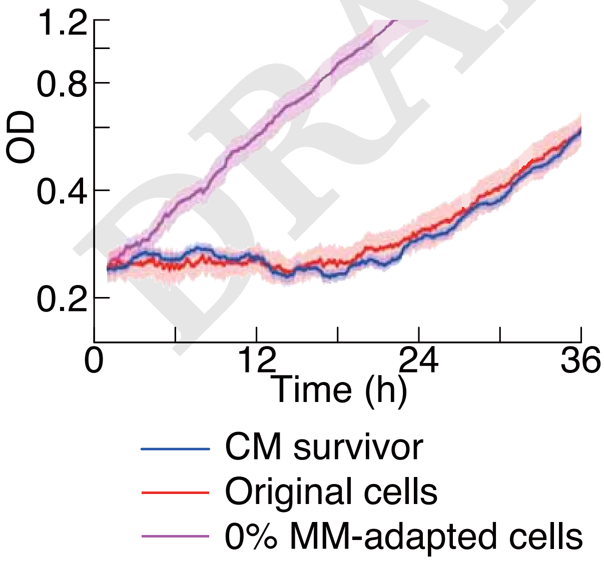


**Supplementary Fig. 2. Growth curves of the cells which survived in WT CM.** The blue line shows the average growth curves of cells in 24 independent colonies isolated from surviving cells in WT CM. Red and magenta lines are the average growth curves of original WT cells precultured in 3% and 0% MM, respectively.

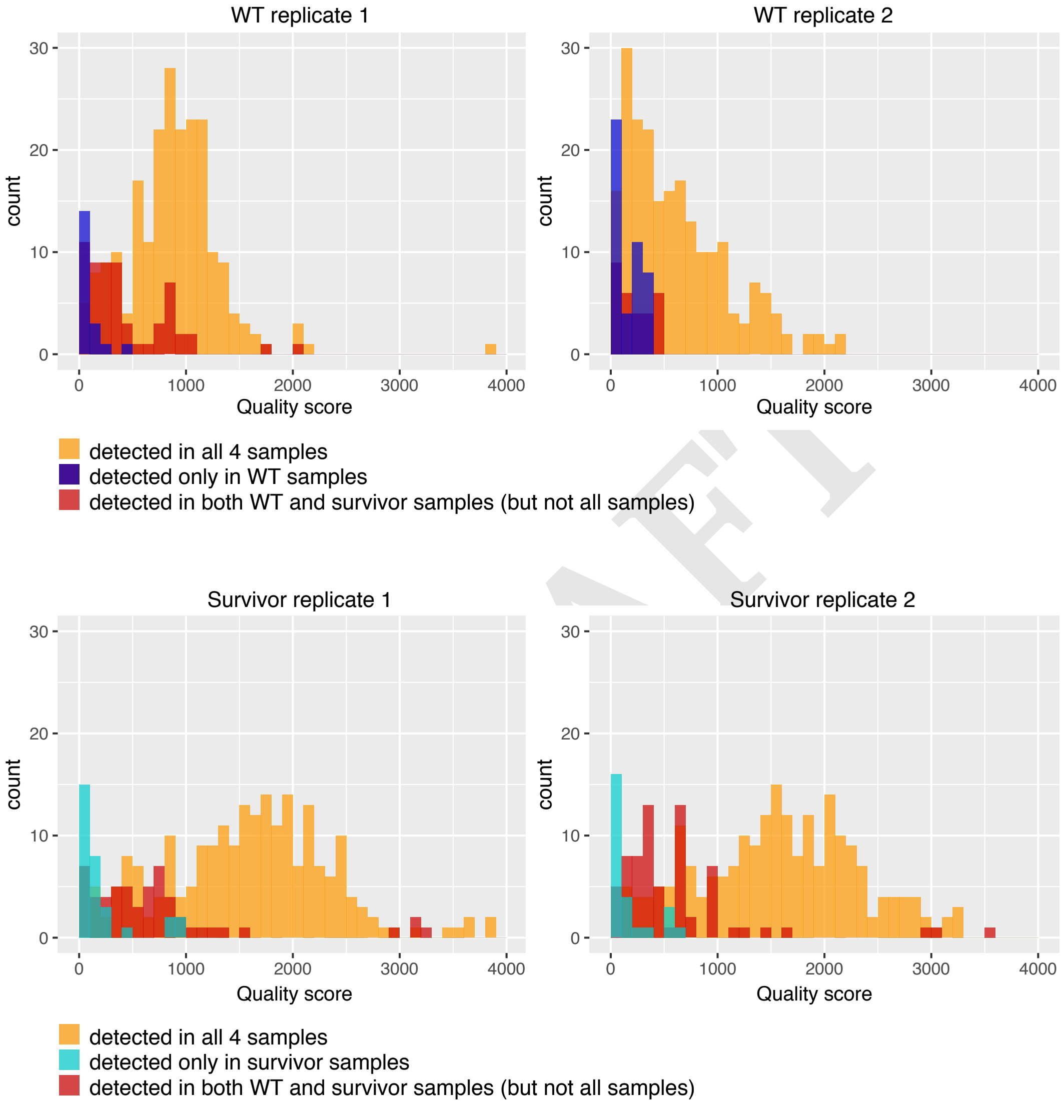


**Supplementary Fig. 3. Histograms of quality scores calculated using GATK for each mutation in original WT cells and cells that survived in WT CM.** The common mutations detected in all four samples are shown in yellow. Mutations detected in both original WT and surviving cells, but not in all four, are shown in red. Unique mutations detected only in a certain replicate of the original WT, and surviving cells are shown in blue and cyan, respectively.

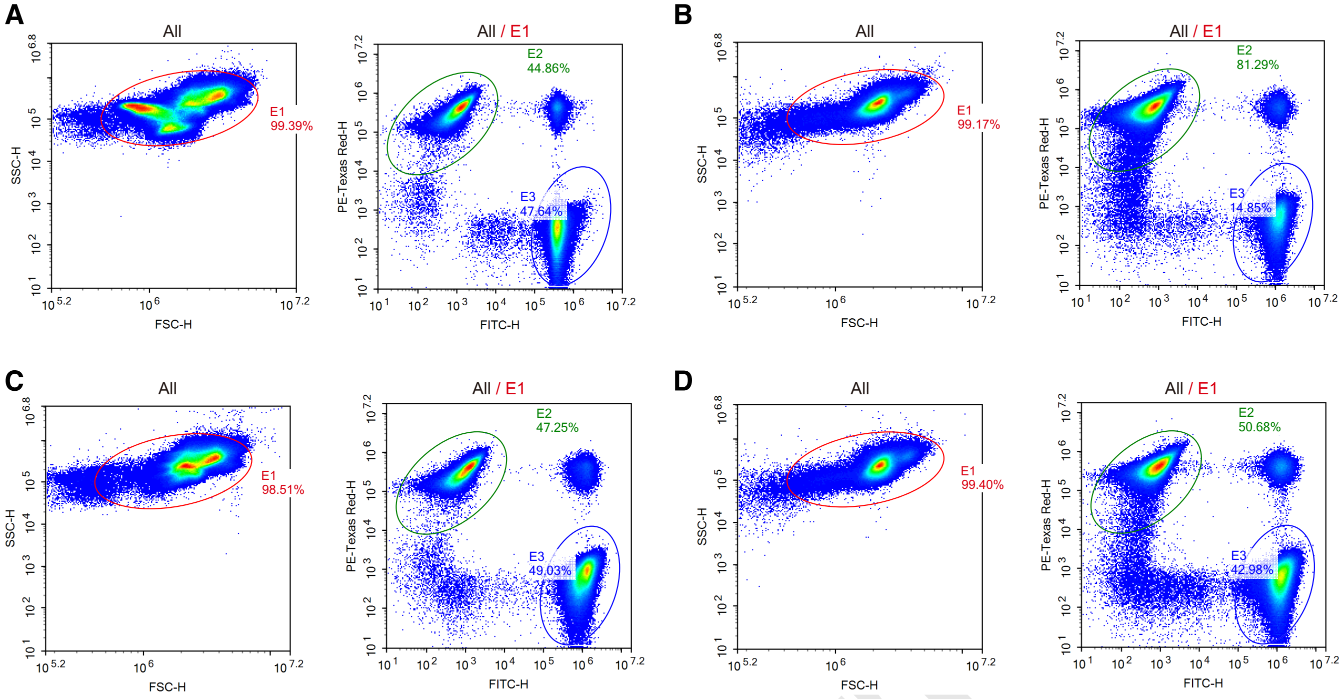


**Supplementary Fig. 4. Flow cytometry gating strategy for identification of the competition assay discussed in Fig.1H and I.** A-D) Fission yeast cells were first gated on the red E1 gate (left panel). Then cells in the E2 area were defined as mCherry positive cells and cells in the E3 area were defined as mNeon-Green positive cells (right panel). The total count of each sample is 500,000 counts. A) An example image of competition assay between mCherry-tagged cells precultured in 0% MM and mNeonGreen-tagged cells precultured in 3% MM at 0h. B) An example image of competition assay between mCherry-tagged cells precultured in 0% MM and mNeonGreen-tagged cells precultured in 3% MM at 72h. C) An example image of competition assay between mCherry-tagged cells precultured in 0% MM and mNeonGreen-tagged cells precultured in 0% MM at 0h. D) An example image of competition assay between mCherry-tagged cells precultured in 0% MM and mNeonGreen-tagged cells precultured in 0% MM at 72h.

DRAFT

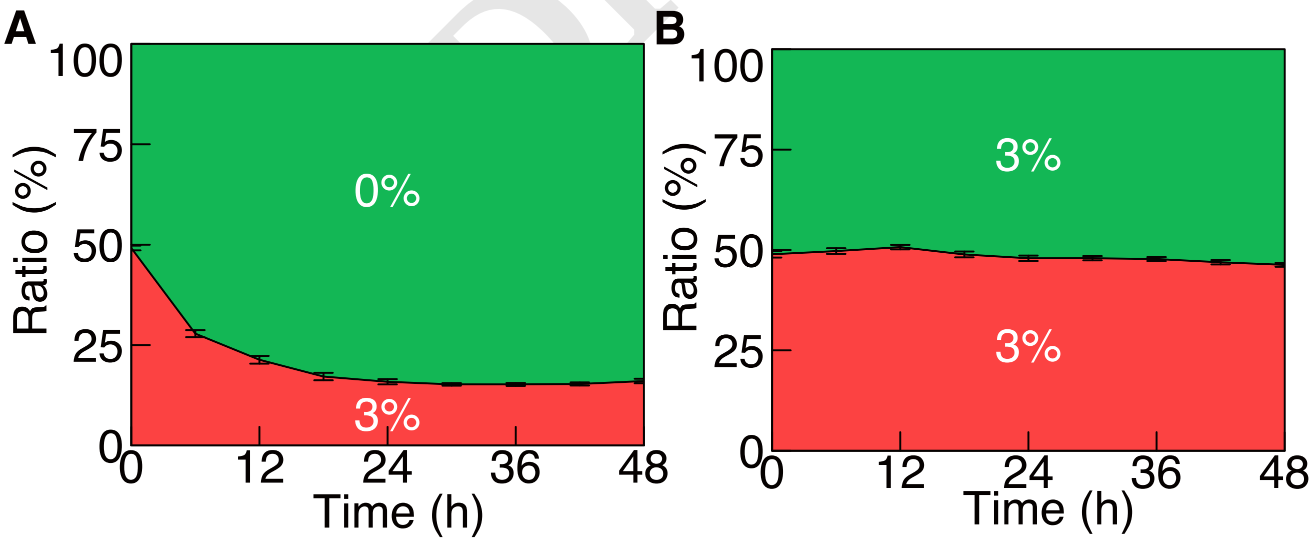


**Supplementary Fig. 5. Release of inhibitory compounds and cell adaptation to them is beneficial for the competition among clonal cells.** mNeonGreen- and mCherry-labelled WT cells were mixed in equal fractions in the WT CM at 0 h, and then they showed population dynamics. Green and red areas indicate the fraction of mNeonGreen- and mCherry-labelled cells, respectively, and overwriting outline characters indicate preculture conditions, that is, 3% and 0% indicate cells precultured in 3% and 0% MM, respectively. Black vertical bars between two areas indicate SEM (number of each sample is 12). (A) Competition assay between mNeonGreen-labelled cells precultured in 0% MM and mCherry-labelled cells precultured in 3% MM. (B) Competition assay between mNeonGreen- and mCherry-labelled cells precultured in 3% MM.

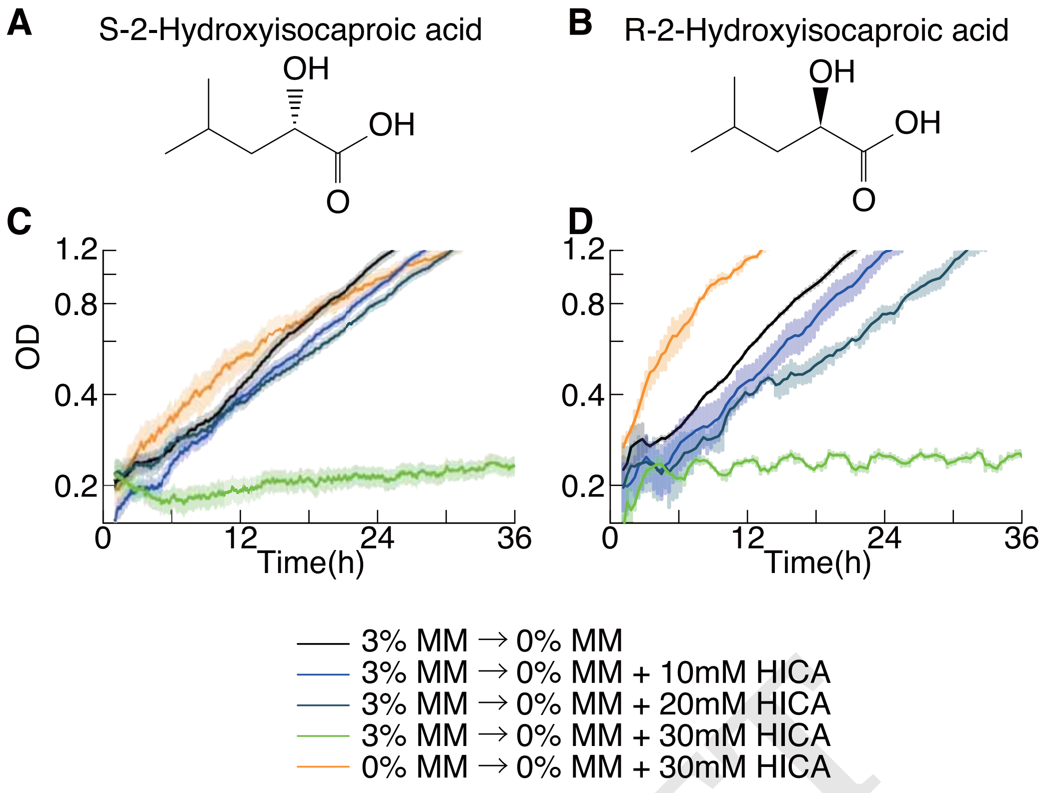


**Supplementary Fig. 6. Growth curves of WT cells in 0% MM with enantiomers of HICA** A and B) Structures of (A) S- and (B) R-forms of HICA. C and D) Growth curves of WT cells in 0% MM with various concentrations of (C) S-form HICA and (D) R-form HICA. For the S-form of HICA, growth curves are an average of 15–18 samples, and for the R-form of HICA, it is an average of two samples.

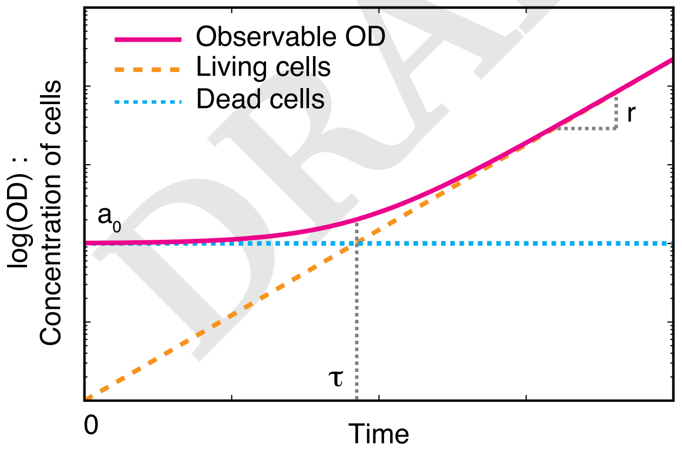


**Supplementary Fig. 7. Schematic representation of growth curves with delay phase.** $a_0$ is the OD at time = 0, $\tau$ is the time when the OD reaches twice of $a_0$, and $r$ is the growth rate at the steady growth phase. The blue dotted line and orange dashed line are the concentrations of dead and living cells, respectively; thus, the solid magenta line, which is the summation of dead and living cells, is observed as OD.

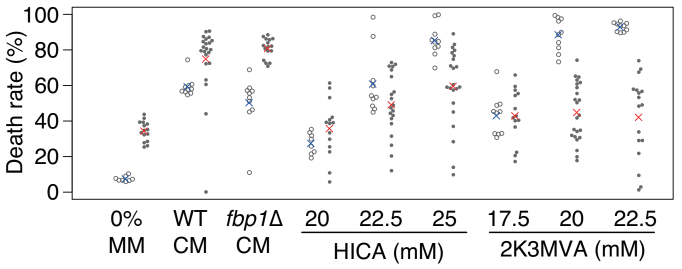


**Supplementary Fig. 8. Measured and estimated death rate in various media.** Open circles represent the death rate measured using phloxine B, as shown in Fig. 5. Blue crosses represent the average of each sample. Filled circles represent the estimated death rate as described in Supplementary Note 2. Red crosses represent the average of each sample.

DRAFT

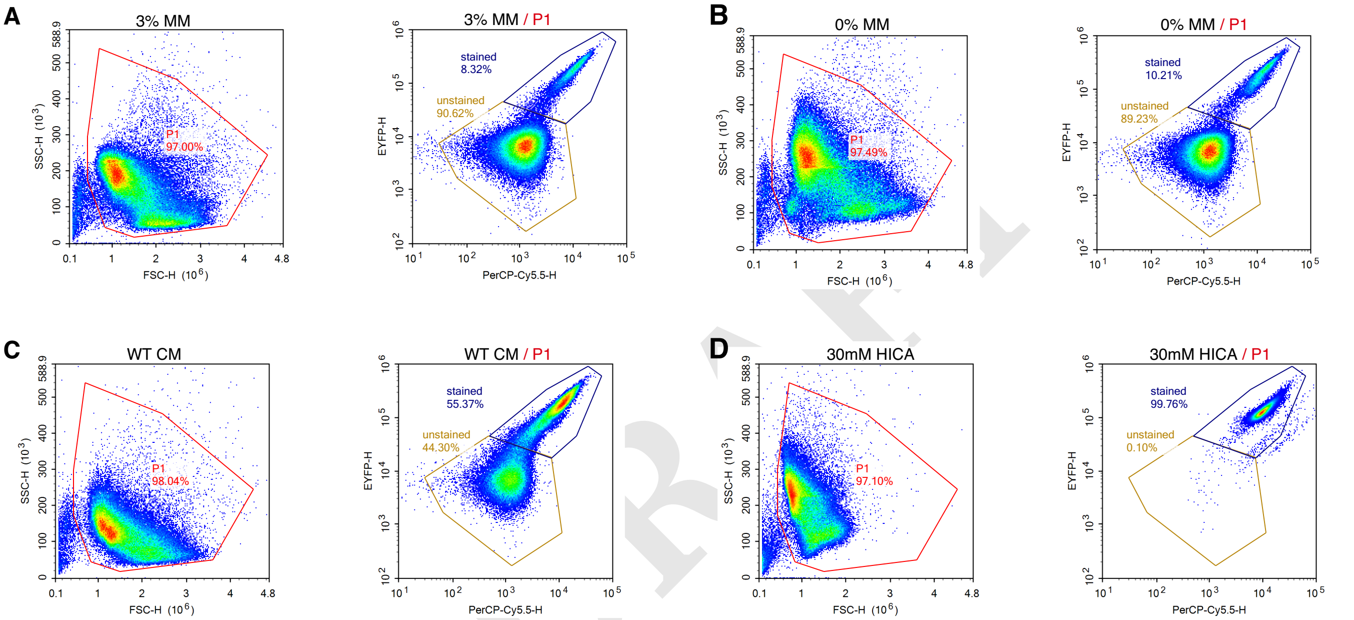


**Supplementary Fig. 9. Flow cytometry gating strategy for identification of the dead cell populations discussed in Fig3G and H.** A-D) Fission yeast cells were first gated on a FSC/SSC scatter plot as the red P1 gate (left panel). Then cells gated on the blue and yellow gates were defined as phloxine B-stained dead cells and unstained living cells, respectively (right panel). The total count of each sample is 100,000 counts. Example images of flow cytometry data for cells in A) 3% MM, B) 0% MM, C) WT CM, and D) 30mM HICA.

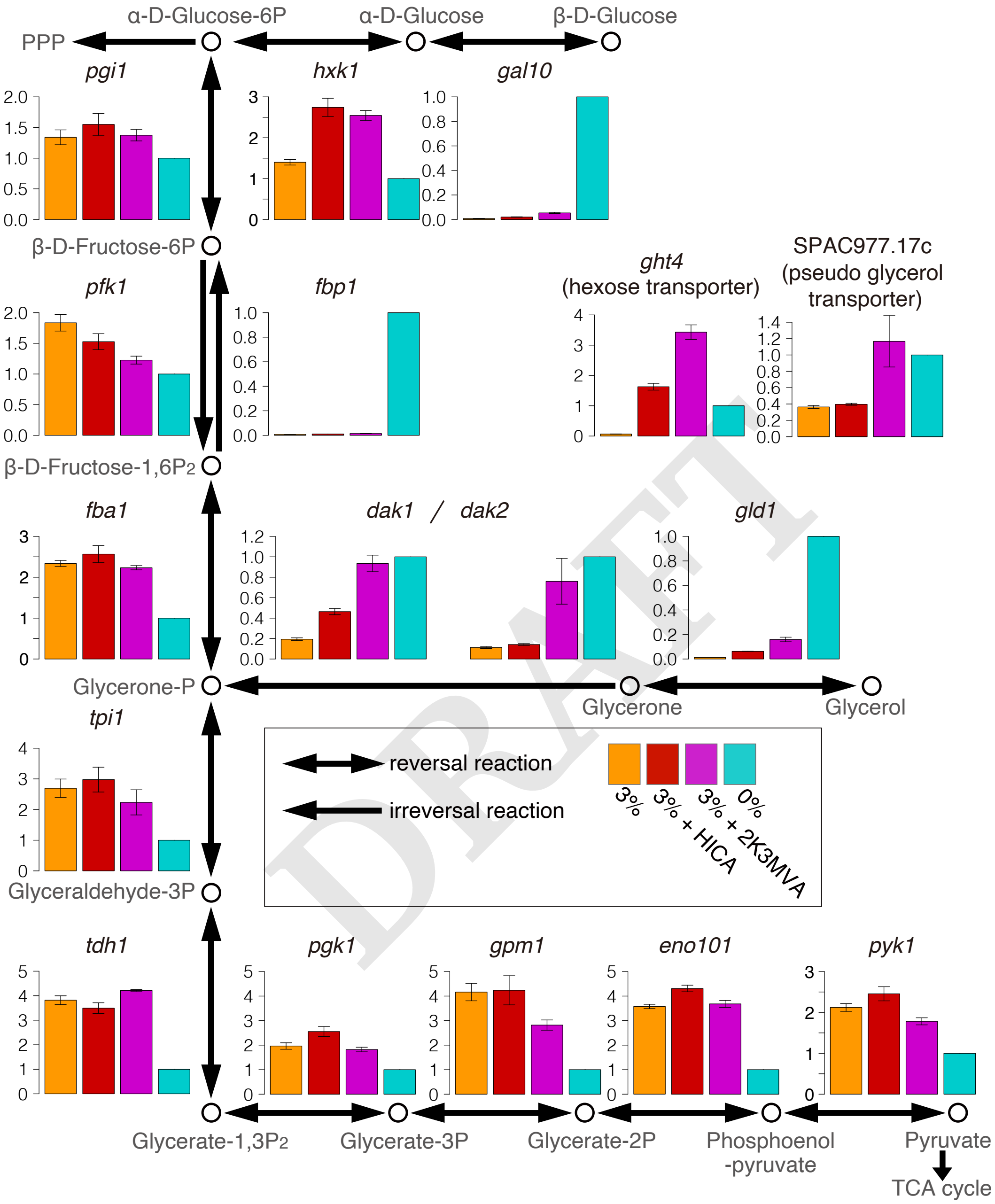


**Supplementary Fig. 10. RNA expression of metabolic genes in 0% and 3% MM with or without the inhibitory molecules.** Expression levels of genes involved in glycolytic and gluconeogenesis pathways in cells cultured in 0% MM (cyan bar), 3% MM (orange bar), 3% MM with 30 mM HICA (red bar), and 3% MM with 25 mM 2K3MVA (purple bar). The gene expression is normalised to that in 0%MM. Vertical bars show SEM, and the sample number of each experiment is 3.

DRAFT

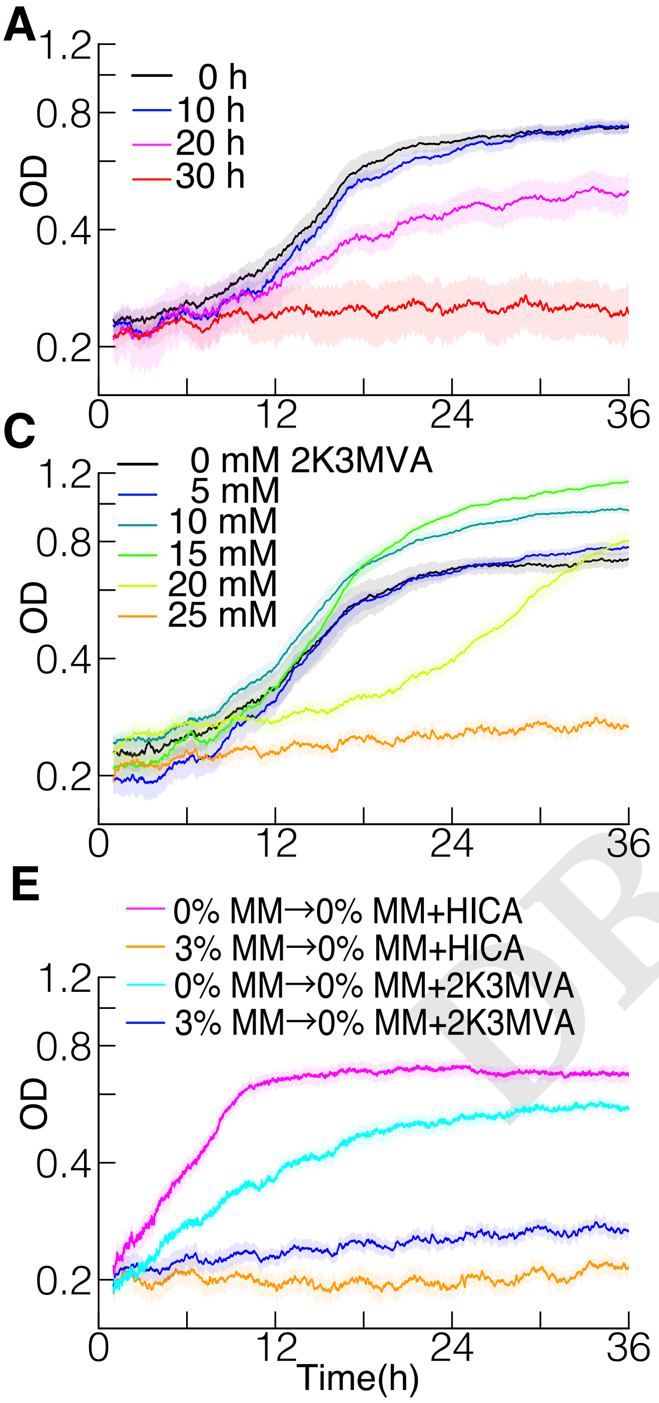


**Supplementary Fig. 11. Media conditioned with various yeast strains caused the delay phase.** A) Growth curves of two strains of *S. cerevisiae* (YEA8) in media conditioned by itself. Different coloured lines indicate growth curves in CM with different incubation times. Each line represents an average of $n \geq 6$ samples. B) Growth curves of YEA8 in 0% MM with various concentrations of HICA. Each line represents an average of $n \geq 5$ samples. C) Growth curves of YEA8 in 0% MM with various concentrations of 2K3MVA. Each line represents an average of $n \geq 5$ samples. D) Growth curves of YEA8 precultured in 0% MM or 3% MM in CM of OC-2. Each line represents an average of $n \geq 6$ samples. E) Growth curves of YEA8 precultured in 0% MM or 3% MM in 0% MM with 25 mM HICA or 25 mM 2K3MVA. Each line represents an average of $n \geq 6$ samples.

### Supplementary Tables

**Supplementary Table 1. List of compounds detected from a minimal medium and conditioned media**
(Abbreviations: N.D., Not detected; N.R., Not reproduced)

| Compound name | Pubchem ID | HMDB ID | m/z | 0% MM | WT CM | *fbp1*Δ *CM* |
|---|---|---|---|---|---|---|
| 2-Hydroxy-4-methylvaleric acid (HICA) | 439960 | HMDB00624 | 131.071 | N.D. | + | + |
| 3-Methyl-2-oxovaleric acid (2K3MVA) | 47 | HMDB0000491 | 129.055 | N.D. | + | + |
| 5-Oxoproline | 7405 | HMDB00267 | 128.035 | N.D. | + | + |
| 2-Hydroxyglutaric acid | 43 | HMDB00606,HMDB00694 | 147.029 | N.D. | + | + |
| Ala | 602 | HMDB00161,HMDB01310 | 90.055 | N.D. | + | + |
| Gln | 738 | HMDB00641,HMDB03423 | 147.076 | N.D. | + | + |
| Glu | 611 | HMDB00148,HMDB03339 | 148.060 | N.D. | + | + |
| Glycerol 3-phosphate (G3P) | 439162 | HMDB00126 | 171.006 | N.D. | + | + |
| Hypoxanthine | 790 | HMDB00157 | 137.045 | N.D. | + | + |
| Inosine | 6021 | HMDB00195 | 269.086 | N.D. | + | + |
| Phe | 994 | HMDB00159 | 166.085 | N.D. | + | + |
| Succinic acid | 1110 | HMDB00254 | 117.019 | N.D. | + | + |
| 1-Methyladenosine | 27476 | HMDB03331 | 282.122 | N.D. | + | N.R. |
| 2-Hydroxyvaleric acid | 98009 | HMDB01863 | 117.055 | N.D. | + | N.R. |
| Adenosine | 60961 | HMDB00050 | 268.104 | N.D. | + | N.R. |
| Citric acid | 311 | HMDB00094 | 191.019 | N.D. | + | N.R. |
| Guanosine | 6802 | HMDB00133 | 284.099 | N.D. | + | N.R. |
| Lactic acid | 612 | HMDB00190,HMDB01311 | 89.024 | N.D. | + | N.D. |
| Pyruvic acid | 1060 | HMDB00243 | 87.009 | N.D. | + | N.D. |
| Tyr | 1153 | HMDB00158 | 182.079 | N.D. | + | N.R. |
| Terephthalic acid | 7489 | HMDB02428 | 165.019 | N.R. | + | + |
| Adenine | 190 | HMDB00034 | 136.061 | + | + | + |
| Glycerol | 753 | HMDB00131 | 93.055 | + | + | + |
| His | 773 | HMDB00177 | 156.077 | + | + | + |
| Leu | 857 | HMDB00687 | 132.102 | + | + | + |
| Pantothenic acid | 6613 | HMDB00210 | 218.104 | + | + | + |
| Uracil | 1174 | HMDB00300 | 113.034 | + | + | + |
| Nicotinic acid | 938 | HMDB01488 | 124.039 | + | + | + |
| Arg | 6322 | HMDB00517,HMDB03416 | 175.119 | N.D. | N.R | + |
| Gly | 750 | HMDB00123 | 76.040 | N.D. | N.R | + |
| Glycerophosphocholine (GPCho) | 439285 | HMDB00086 | 258.109 | N.D. | N.R | + |
| Lys | 866 | HMDB00182,HMDB03405 | 147.112 | N.D. | N.R | + |
| Malic acid | 525 | HMDB00156,HMDB00744 | 133.015 | N.D. | N.R | + |

**Supplementary Table 2. Measured $\tau$, $r$, $a_0$, and estimated $d$**

| Media | $\tau$ (± SEM) | $\tau - \tau_0$ | $r$ (± SEM) | $a_0$ (± SEM) | $d$ (± SEM) |
|---|---|---|---|---|---|
| 0% MM ($n = 15$) | 12.93 (± 0.25) ($\tau_0$) | - | 0.072 (± 0.0018) | 0.25 (± 0.02) | 0.34 (± 0.01) |
| WT CM ($n = 22$) | 32.77 (± 1.03) | 19.84 | 0.056 (± 0.0033) | 0.23 (± 0.01) | 0.75 (± 0.04) |
| *fbp1*Δ CM ($n = 16$) | 33.37 (± 1.31) | 20.44 | 0.057 (± 0.0019) | 0.28 (± 0.02) | 0.81 (± 0.01) |
| 20 mM HICA ($n = 14$) | 14.94 (± 0.78) | 2.01 | 0.065 (± 0.0017) | 0.23 (± 0.02) | 0.36 (± 0.04) |
| 22.5 mM HICA ($n = 20$) | 17.90 (± 0.70) | 4.97 | 0.064 (± 0.0019) | 0.23 (± 0.02) | 0.49 (± 0.04) |
| 25 mM HICA ($n = 22$) | 29.63 (± 1.62) | 16.70 | 0.047 (± 0.0020) | 0.21 (± 0.02) | 0.60 (± 0.05) |
| 17.5 mM 2K3MVA ($n = 14$) | 17.45 (± 1.23) | 4.52 | 0.061 (± 0.0028) | 0.22 (± 0.02) | 0.43 (± 0.04) |
| 20 mM 2K3MVA ($n = 23$) | 22.98 (± 1.19) | 10.06 | 0.048 (± 0.0020) | 0.25 (± 0.03) | 0.45 (± 0.03) |
| 22.5 mM 2K3MVA ($n = 17$) | 36.55 (± 3.16) | 23.63 | 0.031 (± 0.0018) | 0.24 (± 0.02) | 0.42 (± 0.06) |

DRAFT

**Supplementary Table 3. Strains used in this study**

| Organism | Strain name | Genotype | Description | Reference |
|---|---|---|---|---|
| *S. pombe* | L972 | $h^-$ | Wile type (WT) | (10) |
| *S. pombe* | AR110 | $h^-$ deletion of Chr2:197561-200480 | *fbp1*$\Delta$ | this study |
| *S. pombe* | HN98 | $h^-$ SPBC1348.11<<KanMX6-Padh1-mCherry | mCherry-tagged | this study |
| *S. pombe* | HN101 | $h^-$ SPBC1348.11<<KanMX6-Padh1-mNeonGreen | mNeonGreen-tagged | this study |
| *S. cerevisiae* | OC-2 | Wild type | Wine yeast IAM4274, JCM1419 | (11) |
| *S. cerevisiae* | YEA8 | Wild type | Baker's yeast, Lab stock | this study |

**Supplementary Table 4. PCR primers for RT-PCR used in this study**

(Abbreviations: FW, forward primer; RV, reverse primer)

| Target genes | FW/RV | Sequence (5'-3') |
|---|---|---|
| *fbp1* | FW | TCGCCAACACCATTCGTAAAGC |
| *fbp1* | RV | ACGATGAGTTTGCAGCATCCGTTCG |
| *tdh1* | FW | CGCTTCGATGGCTCCGTCGAG |
| *tdh1* | RV | CACGCTTGGCACCACCCTTC |
| *eno101* | FW | AAGGGTCCCTATGTCCTCCC |
| *eno101* | RV | GTGTGGTAGGTCTCAGCACC |
| *ght4* | FW | GTGGTGCAGTTGTTTCGACC |
| *ght4* | RV | AGGTAACGCGGTGATTCAGG |
| *gld1* | FW | GCCTCCTCTGATGCCGCTAC |
| *gld1* | RV | CCACACCTCCGGCAAAGGAA |
| *dak1* | FW | ATCGGTGCTTCACTCGCTCA |
| *dak1* | RV | GGCACGGTCCTTGTCGGATT |
| *pgi1* | FW | AGACCGTGCCGTCTTTCACC |
| *pgi1* | RV | ACCATAACGGGGCCCAAGTC |
| *tpi1* | FW | TGGTGAGACTTTGGCCGAGC |
| *tpi1* | RV | GTTGGTAGCCCACTTGCGGA |
| SPAC977.17 | FW | CAACCCAGGGCATGACTGCT |
| SPAC977.17 | RV | CCTCCTATGGTACCGCCCCA |
| *pyk1* | FW | ACCCCGTTGAAGCCGTTACC |
| *pyk1* | RV | GCAGTGTTACCGCTGGTGGA |
| *gpm1* | FW | AGTCCAAATCTGGCGCCGTT |
| *gpm1* | RV | GGACACCAGTGGCAAGCTCA |
| *fba1* | FW | TGCTGCTCTTGAGGCTGCTC |
| *fba1* | RV | CTTGGCGCAGTGGTCAGAGT |
| *pfk1* | FW | CCCGCTGCCGAAGATTCTGA |
| *pfk1* | RV | AGCACCAGACTCCTCGCTCT |
| *dak2* | FW | TCTGGTGGTGGTAGTGGGCA |
| *dak2* | RV | CTCTCAGCTGCCAGGCCAAA |
| *hxk1* | FW | CCCCTGTGTTCAACGCTCCA |
| *hxk1* | RV | TCTCAGTACCTGGGCTGGCA |
| *gal10* | FW | TCCAATCCCAGAGTCGTGTCCT |
| *gal10* | RV | CTCGCCTACCGATGGCTACTTG |
| *pgk1* | FW | GCCCGTATCGTTGGTGCTCT |
| *pgk1* | RV | CCTCACCACCCTTGGCTTCC |

DRAFT


DRAFT
